# Monitoring Alzheimer’s Disease via Ultraweak Photon Emission

**DOI:** 10.1101/2023.03.14.532685

**Authors:** Niloofar Sefati, Tahereh Esmaeilpour, Vahid Salari, Asadollah Zarifkar, Farzaneh Dehghani, Mahdi Khorsand Ghaffari, Noémi Császár, István Bókkon, Serafim Rodrigues, Daniel Oblak

## Abstract

The present study takes on an innovative experiment involving detection of ultraweak photon emission (UPE) from the hippocampus of male rat brains and finds significant correlations between Alzheimer’s disease (AD), memory decline, oxidative stress, and the intensity of UPE emitted spontaneously from the hippocampus. These remarkable findings opens up novel methods for screening, detecting, diagnosing and classifying neurodegenerative diseases (and associated sydromes), such as in AD. This also paves the way towards novel advanced brain-computer interfaces (BCIs) photonic chip for the detection of UPE from brain’s neural tissue. The envisaged BCI photonic chip (BCIPC) would be minimally invasive, cheap, high-speed, scalable, would provide high spatiotemporal resolution of brain’s activity and would provide short- and long-term screening of clinical patho-neurophysiological signatures, which could be monitored by a smart wristwatch or smartphone via a wireless connection.

**Background & aim:** Living cells spontaneously emit biophotons, or UPE, during the process of metabolic reactions, and these UPE in tissues may be altered in pathological conditions. These compelling observations led us to hypothesise that AD (a severe neuropathological disorder) can be screened via UPE. This is substantiated by previous studies showing that oxidative stress occurs prior to the formation of amyloid plaques and neurofibrillary tangles (i.e. the neuropathological hallmarks of AD). Indeed, oxidative stress is a critical factor contributing to the initiation and progression of AD. Moreover, earlier research have evidenced the association between UPE and oxidative stress of biological tissue. These combined observations set us to investigate whether UPE intensity of the hippocampus in a pathological state, induced by intracerebroventricular (ICV) injection of streptozotocin (STZ), can be correlated with memory, oxidative stress, Acetylcholinesterase (AChE) as a novel screening strategy for AD.

**Material & methods:** Thirty-two adult male rats were divided into four groups: Control, Sham, STZ, and STZ+Donp (n=8). Specifically, for inducing sporadic AD (sAD), STZ was injected on days 1 and 3. One week after the second ICV injection, the intraperitoneal (IP) use of donepezil was initiated and continued for two weeks. After treatment, spatial and recognition memory were evaluated from days 24 to 29 of the experiment using the Morris water maze (MWM) and novel object recognition (NOR) test, respectively. Finally, the rats were euthanased by cervical dislocate in day 30. Anesthetic drugs disrupt neural communication from chemical neurotransmitter receptor inhibition. UPE related to cells activity so anesthesia intervention must be considered. Then, their brains were removed and the hippocampus dissected. The Right hippocampus was evaluated in terms of UPE via a Photomultiplier tubes (PMT) device. Moreover, in left hippocampus we measured malondialdehyde (MDA) by the TBARS assay and heat via calorimeter ELIZA device. Acetylcholinesterase (AChE) activity was also scrutinized via acetylthiocholine reaction via the Ellman method.

**Results & discussion:** STZ injection impaired learning and memory function compared with the sham and control groups. The results of the MWM test indicated a decrease in the time used to find the hidden platform in the donepezil-treated group during training days, while in the STZ group, no significant reduction in this time was observed. In the probe trial, the donepezil-treated group showed a significant increase in target quadrant time in comparison with the STZ group (p<0.05). Furthermore, the object recognition test demonstrated that the donepezil-treated group spent more time recognizing new objects in the testing phase (p<0.05). Whereas, in the STZ group, there was no significant difference in spent time for identifying the objects. Ex vivo detection of UPE from the hippocampus of rats showed that the sham group had higher UPE than the Control group (p<0.05). The STZ injection significantly increased UPE and MDA concentrations in the hippocampus than in the Sham and Control groups (p<0.0001). Correlation analysis of results reveal that the emission intensity is associated with the MDA concentration (r = 0.855). Hippocampus AChE activity also significantly increased in STZ-injected groups. Treatment with donepezil decreased MDA concentration, UPE intensity, and activity of AChE in comparison with the STZ group (p<0.05). UPE intensity was linked with AChE activity as evidenced by Pearson correlation analysis between UPE intensity and AChE activity (r = 0.779). Conclusion: The hippocampus UPE increases in STZ-induced sAD and is associated with the redox state of the tissue. Donepezil decreases the UPE and improves the oxidative stress induced by STZ injection. Since oxidative stress is one of the primary hallmarks in the progression of AD, then it stands to reason that the Brain’s UPE emission can be used as a novel methodology for screening AD. Moreover, UPE could be used to monitor recovery from neurodegenerative diseases upon suitable future therapeutic treatments, as suggested by our experiment involving donepezil. Our findings, encourages further research and suggests the development of a minimally invasive BCI photonic chip (with similar quantum efficiency as PMT) for screening and diagnosing AD.

## 1 INTRODUCTION

To the best of our knowledge, there is currently no brain implantable chip that is specifically designed to screen and diagnose Alzheimer’s disease (AD). While implantable brain chips have been developed for various purposes, such as monitoring brain activity or delivering therapeutic electrical stimulation, their use for screening or diagnosing AD is still an area of active research. Diagnosing AD typically involves a combination of cognitive and memory tests, brain imaging studies, and other assessments performed by a clinician. Based on the observation that the brain spontaneously emits photons, so called ultraweak photon emissions (UPE) (1; 2), we suggest that a brain-computer interface with an integrated photonic chip (BCIPC)(3) may be an efficient real-time method for monitoring early symptoms of AD and related dementias (ADRD). The envisaged technology would support clinicians by providing complementary data to efficiently screen and diagnose AD. Indeed, We have previously discussed a pattern recognition approach for an efficient interpretation of UPE via the output signals on a photonic interferometer chip (3). Supporting this vision is the fact that various research studies have shown that there are significant UPEs from neurons across the electromagnetic spectrum (1; 4). Such UPEs reflect cellular (and brain) oxidative status, as they are particularly intense during heightened metabolic activity or stress (5). Several studies point to direct correlations between UPE intensity and neural activity, oxidative reactions, EEG activity, cerebral blood flow, cerebral energy metabolism, and glutamate release (6). Such correlations suggest that we may use UPEs as a correlative signal to monitor different internal states across the stages of ADRD pathology and to expand the clinical criteria, particularly in the preclinical and mild cognitive impairment (MCI) stages where memory loss and other problems are not always evident. Discrimination between the interferometric patterns of normal, and preclinical stages will be non-trivial but tractable via machine learning, based on observation of highly synchronized brain activities with strong UPE correlations for specific cognitive tasks (3). With an analysis of signals over thousands of training trials, it will be possible to obtain an average pattern for feature extraction, enabling pattern recognition directly during preclinical and premarket approval testing.

The hippocampus is an important brain region that plays a crucial role in forming and retrieving memories (7). In AD, one of the earliest symptoms is memory loss and difficulty in forming new memories, which is linked to damage to the hippocampus. This is why the hippocampus is often a focus of research and imaging studies in the diagnosis of AD. The size of the hippocampus can also decrease in people with AD, which can be seen on brain scans such as MRI. However, it’s important to note that memory loss and changes in the hippocampus can be caused by other factors as well, so it’s not a definitive diagnostic tool for AD, hence a number of clinical evaluations are undertaken to accurately diagnose AD.

The detection of biophotons through a photonic chip implanted in the hippocampus is an area of ongoing research, and its potential for diagnosing AD is still unclear. Biophotons are extremely weak light emissions from biological systems and their relationship to neurodegenerative diseases such as AD is not yet fully understood, and there is currently no evidence to support its use as a reliable diagnostic tool for AD. This motivates the present study where we investigate biophotons from the hippocampus (8) and correlate it with other neuropahtological signatures of AD, results of which enables us to propose the future development of BCIPC for screening and diagnosing AD.

### 1.1 Alzheimer’s disease (AD), ROS, and Ultraweak Photon Emission

AD is the most common type of dementia and is a progressive neurodegenerative brain disorder causing a significant disruption of normal brain structure and function (9). Sporadic Alzheimer’s disease (sAD), which begins after the age of 65 without a family history, is the most common type of AD and has several causes and risk factors (10). The risk factors and reasons associated with sAD include the accumulation of amyloid plaques, the formation of neurofibrillary tangles, decreased activity or number of cholinergic neurons in the brain, neuroinflammation, insulin signaling impairments, mitochondrial disorders, and oxidative stress (11; 12). In the brain of an AD patient, the most consistent neurotransmitter-related change is the reduction of cholinergic innervation in the cortex and hippocampus caused by the loss of neurons in the basal forebrain (13). Among the pharmacological agents, Acetylcholinesterase (AChE) inhibitors, like donepezil, seem to be the most effective agent for improving cholinergic deficits and reducing the symptoms of AD (14). As the most potent approved drug, donepezil affects various events of AD, such as inhibiting cholinesterase activities, anti-Aβ aggregation, anti-oxidative stress, etc. (15). In sAD, the initiating causes of the neurodegenerative cascade are unknown, but some studies suggest increased levels of oxidative stress and impaired energy metabolism as the initiating cause of the disease (16). Oxidative stress is an imbalance between the antioxidant defense system and the production of reactive oxygen species (ROS). Mitochondria are susceptible to oxidative damage despite the presence of an antioxidant system, and damaged mitochondria produce more ROS than ATP (17). Spontaneously, when ROS are produced during the metabolic processes, the ultra-weak photons are emitted through the relaxation of electronically excited species formed during the oxidative metabolic processes (18); therefore, the biophoton emission rate could be utilized in order to investigate tissue oxidative state (19). The ultraweak photon emission (UPE) produces a very weak luminescence and can be performed by living organisms (18; 20), comprising microorganisms, plants, and humans. It is mainly named biophoton emission (21; 22). The UPE intensity changes are related to different physiological and pathological conditions, such as different kinds of stress, mitochondrial respiratory chain, cell cycle, and cancerous growth. It has been shown that the measurement of delayed luminescence emitted from the tissues provides valid and predictive information about the functional status of biological systems (23; 24). Several studies repeatedly illustrated that the intensity of photon emission changes in an abnormal condition, and abnormal cells emit significantly more biophotons than healthy cells. It has also been shown that changes in biophotonic activity are indicative of changes in mitochondrial ATP energy production manifested in physiological and pathological conditions (25; 26). Having considered that ROS production is related to inflammatory diseases and impaired metabolic processes (27), it is reasonable to expect that UPE can also be associated with inflammatory disease and/or metabolic processes (28). Therefore, UPE might be used practically for the diagnosis of inflammation and inflammation-related diseases (28). Reports have considered ultra-weak photon emission as a potential diagnostic tool, and some studies have found evidence for diagnosing patients with type 2 diabetes (29) and breast cancer (30). Intracerebroventricular (ICV) injection of low doses of streptozotocin (STZ) leads to neuropathological, biochemical, and behavioral changes similar to non-hereditary sAD in the rat brain, so it is used as a laboratory model to investigate the process and treatment of sAD in rat brain (31). STZ possibly desensitizes neuronal insulin receptors and reduces the activities of the glycolytic enzyme (32). It causes oxidative stress and decreases cerebral energy metabolism resulting in cognitive dysfunction by inhibiting the synthesis of adenosine triphosphate (ATP) and acetyl CoA, which in turn leads to cholinergic deficiency supported by reduced choline acetyltransferase (ChAT) activity in the hippocampus (33; 34) and increased AChE activity in rat whole brain (35). The present study was designed to evaluate the UPE intensity of the hippocampus in the normal, pathological, and therapeutical state and also investigate the relationships between the intensity of UPE, memory, oxidative stress, and AChE activity for AD diagnosis and treatment success.

## 2 MATERIALS AND METHODS

The whole method is graphically represented in Fig.1. The details are as follows:

**Figure 1.**
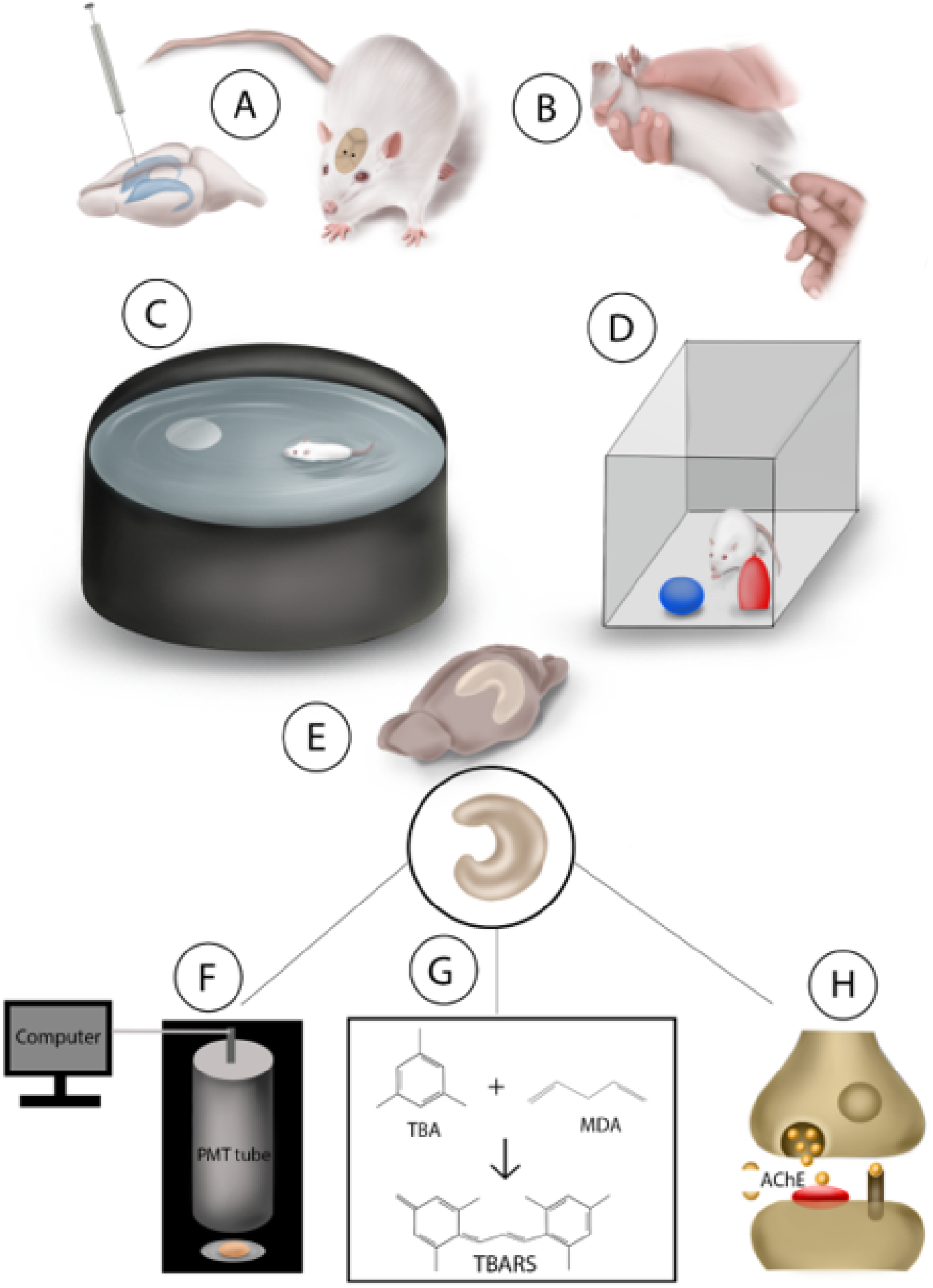
(A), ICV-STZ Injection; (B), IP injection of treatments; (C), Morris Water Maze test;(D), Novel Object Recognition test; (E), Hippocampus dissection; (F), UPE detecting of the right hippocampus with photon multiplier tube; (G), MDA estimation of left hippocampus by colorimetrical measurement of TBARS reaction with ELASA reader; (H), measurement of AChE activity of the left hippocampus with Ellman method

### 2.1 Animals

In this study, 32 adult Sprague-Dawley male rats (220-250 g), were obtained from the Comparative and Experimental Medical Center of the Shiraz University of Medical Sciences. All rats were housed under standard conditions (temperature: 22±2 °c, relative humidity: 50%, with a 12-hour light/dark cycle) and had free access to laboratory food and water. The animals were subjected to an acclimatization period of one week before the beginning of the experiments. To prevent seasonal disturbances, experiments were started on all groups at the same time. All procedures in the study were based on the National Institutes of Health (NIH) guidelines for the care and use of laboratory animals (NIH Publications No. 8023, revised 1978) and were approved by the Ethics Committee of the Shiraz University of Medical Sciences (approval number: IR.SUMS.REC.1400.191).

### 2.2 Drugs and reagents

Donepezil, was produced by Sigma, St. Louis, USA, and STZ, thiobarbituric acid, and trichloroacetic acid were procured from the Sigma-Aldrich company. Artificial cerebrospinal fluid (aCSF) was prepared as follows: (in mmol/L) 147 mM NaCl, 2.9 mM KCl, 1.6 mM MgCl2, 1.7 mM CaCl2, and 2.2 mM dextrose.

### 2.3 Experimental design

The rats were divided with simple randomized sampling into four groups (n=8), including Control, Sham, STZ, and STZ+Donp groups. the Control group underwent without any intervention. Sham and experimental groups underwent ICV cannulation of both lateral ventricles. To produce the ICV-STZ rat model and memory impairment in rats, after cannulation, ICV injection of STZ on days 1 and 3 of the experiment was done. For this purpose, 3 mg/kg STZ (Sigma, St. Louis, USA) was dissolved in 10 *μ*l sterile saline 0.9% and injected slowly into both lateral ventricles (each side 5μl). The Sham rats were treated identically with 10 μl sterile saline 0.9%. One week after the second injection of STZ, the intraperitoneal (IP) treatments were initiated on the rats until day 23. Sham and STZ group received 0.2 ml saline 0.9% daily/IP, and the STZ+Donp group received 0.75 mg/kg donepezil daily/IP.

From days 24 to 29 of the experiment, rats were subjected to MWM and NOR tests. Finally, on day 30, the rats were dislocated, and their brains were removed (see Fig.2).

**Figure 2.**
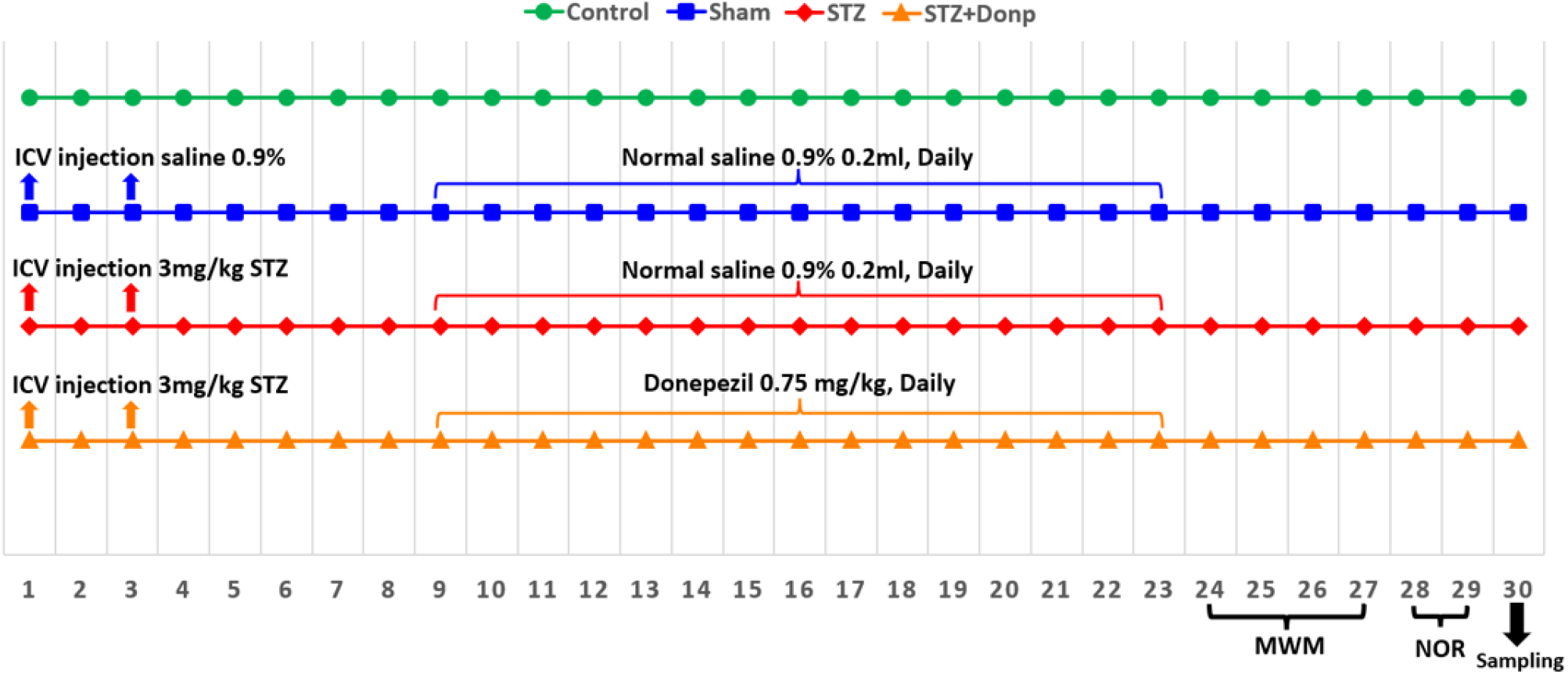
Timeline of experiment. Lines represents groups.

### 2.4 Intracerebroventricular cannulation

On the zero-day of the experiment, we carried out ICV cannulation of both lateral ventricles. Briefly, rats were anesthetized with an IP injection of ketamine HCl 10% (70 mg/kg) and xylazine HCl 2% (10 mg/kg). Their heads were placed in a stereotaxic frame, and a midline incision was made sagittally in the scalp. The bregma boundary was visible after removing the remaining tissue. Holes were drilled in the skull with a dental handpiece with a burr size of 1mm on both sides over the lateral ventricles using the following coordinates from Paxinos atlas (0.8 mm posterior to bregma, 1.5 mm lateral to the sagittal line). Two cannulas with a height of 3 mm were inserted in these holes. Then, holes and cannula were covered with dental cement to fix the position. For ICV injection, the injection needle with 3.5 mm height was connected to the 10 *μ*l Hamilton syringe (Bonaduz, Switzerland) by a short piece of narrow polyethylene tube. The needle was inserted into the tip of the cannula.

### 2.5 Morris water maze

MWM test was performed from days 24 to 27 of the experiment. The maze consisted of a circular pool (160 cm in diameter, 60 cm in height), and up to 35 cm in height, it is filled with water (the temperature at 22 ± 1°C). The pool is geographically divided into four quarters, equal north, south, east, and west, and in each quarter of the circle, a point is intended to leave the animal in the water. A transparent platform (diameter, 10 cm) was positioned in the middle of the target quadrant and submerged approximately 1 cm below the water’s surface. All rats underwent four daily trials for three consecutive days for spatial learning assessment. The starting quadrant was changed each day. If the rat failed to find the hidden platform within 60 s, it was guided to the platform by the experimenter. The rats stood on the platform for 10 s for spatial examination of the platform zone. Then, they were removed from the pool into the cage and rested for one min under a heater inside the cage. A probe trial was performed 24 h after the last acquisition trial to assess spatial memory. In this phase, the hidden platform was removed, and rats were abandoned from the opposite quadrant of the target quadrant into the water and given 60 s to swim in the pool. A visible platform trial was performed after the probe trial to check the rat’s vision and platform perception. Time to reach the platform (escape latency), time spent in the target quadrant, the number of platform site crossings, and swimming speeds were automatically estimated with a video tracking system (EthoVision XT, Noldus Information Technology) for measuring mobility accurately; the software must be calibrated.

### 2.6 Novel object recognition test

On the 28th day of the experiment, recognition memory was evaluated by a novel object recognition test. The testing apparatus was a box with dimensions 65× 45 × 65 cm. this test was performed for two days. On the first day, to familiarize themselves with the test box, the rats are located in the test box for 5 min without any objects. On the second day in the familiarization phase, two similar objects were placed in two corners of the box, and the rats were placed in the box for 5 min to explore objects. Then the rats returned to the cage. After 60 min, the rats were retrial in the box for the testing phase, and one of the familiar objects was replaced with a new object. The time spent to check each object was measured for 5 min. The animals were evaluated when facing, sniffing, or biting the object. some equipment, such as a test box and objects, were cleaned with 70% ethanol between trials. Eventually, the discrimination index (the spent time identifying a novel object divided by the spent total time exploring either object) was measured for recognition memory assessment.

### 2.7 Hippocampus sampling

Finally, the rats cevical dislocated on day 30, and the brains were rapidly removed. Then, the right and left brain hemispheres separated, the right hippocampus was dissected for biophoton emission evaluations, and the left one was stored at −80°C for MDA and AChE activity measurements (see Fig.3).

**Figure 3.**
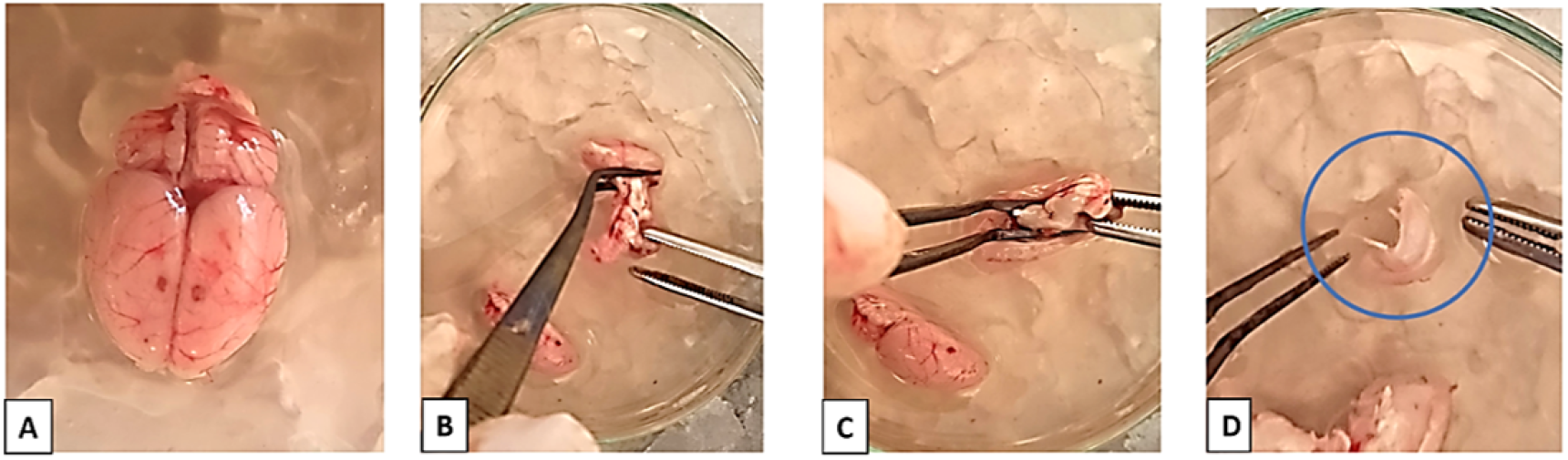
Hippocampus dissection. (A) The brain was dissected, (B) Hemisphere was separated, and the cerebellum, pons, and medulla were removed, (C) Thalamus and hypothalamus dissection, (D) Hippocampus was removed from the cortex and placed in a petri dish containing ACSF.

### 2.8 Detection of UPE

In this study, UPE were detected with the photomultiplier tube (PMT) placed in a dark box in a dark room. PMT is an intensely sensitive detector amplifying entrance photons from a field of view to electrical signals. The PMT was connected to the G.G.104 (Parto-Tajhiz-Besat co - PTB) converter, which was also connected to the laptop for data to be digitally visible. A photon counting system (R6095 Hamamatsu Photonics K.K., Electron Tube Center, Hamamatsu, Japan) was used to observe time-dependent photon emission intensity. PMT provides detection of photons in the range of 300 to 700 nm wavelength, with highest quantum efficiency (30%) at 420 nm. The collecting gate time from the PMT was set at 1s. Dark noise was detected with the number of counts in an empty dark box for 5min (c.p.5 min) before sample UPE detection and subtracted from the results. Noise is reduced by modifying the upper and lower threshold via PMT software. The distance between the sample and the PMT sensor was 0.5 cm. In each trial period, the medium’s emission and then the UPE of samples were measured in a 5 min period. The right hippocampus was dissected and transferred to a 3 cm Petri dish containing oxygenated aCSF (O_2_ 95%, CO_2_ 5%), which was placed under the sensor. For declining any possible delayed luminescence, petri dish was placed in the darkroom for 10 min (23).

### 2.9 Sample preparation for biochemical analysis

The left hippocampus was removed, weighed, and homogenized in the 10-fold ice-cold phosphate buffer saline. The homogenizing was accomplished using Homogenizer (IKA T10 basic, Germany) apparatus for about 3 m. Next, centrifuging (12000 rpm) at 4°C for 5 m was performed, and the supernatant was isolated for the following assessments.

### 2.10 MDA Assessment

MDA, a marker of lipid peroxidation, was estimated colorimetrically using 1,1,3,3-tetra ethoxy propane as a standard. Lipid peroxidation was estimated with TBARS and determined by colorimetric measurement of the color produced during the reaction of thiobarbituric acid (TBA) with MDA by an Elisa plate reader (Bio-Tek Instruments, Inc) at 532nm (37). Results expressed as nmol/mg protein.

### 2.11 Acetylcholinesterase activity assessment

The activity of AChE was carried out according to the Ellman method, which uses acetylthiocholine as a substrate. Thiocholine, produced by AChE, reacts with 5,5-dithiobis (2-nitrobenzoic acid) to form a colorimetric product proportional to the AChE activity. The activity of AChE was spectrophotometrically measured at 412 nm and expressed as nmol/min/mg protein (38). Protein concentration was measured according to the method of Bradford (39).

### 2.12 Statistical analysis

Data were presented as the mean ± SEM, and P < 0.05 was considered statistically significant. To assess escape latency and swim velocity (Morris water maze) changes during the time, a two-way repeated-measures analysis of variance (TWRM-ANOVA) was done. Also, for other parameters, one-way ANOVA followed by Tukey’s post hoc test was utilized for comparing various groups. In the NOR test, exploration time during the testing phase was analyzed by paired t-test. All statistical analyses were performed using GraphPad software (Prism Software Inc., San Diego, CA, USA).

## 3 RESULTS

### 3.1 Morris Water Maze

There was no significant difference in escape latency between groups on the first day of the acquisition trial. On the 2nd and 3rd days, there was a significant difference in escape latency between the control group and other groups (STZ p < 0.0001, STZ+Donp p < 0.05). STZ group had significantly higher scape latency in comparison with the sham group (p < 0.0001). Also, on this day, the escape latency of the STZ+Donp group significantly decreased in comparison with the STZ group (p < 0.001). Our results demonstrated that escape latency in the donepezil-treated group was significantly lower than in the STZ group (p < 0.001) (see Fig.4A). No significant differences were observed in swim speed between groups on all days of the trial (Fig.4B). In the probe trial, the STZ-injected groups significantly spent lower time in the target quadrant compared with the control group (STZ p < 0.0001, STZ+Donp p < 0.01). Still, only the STZ group was significant with the sham group (p < 0.001). Whereas the donepezil-treated group significantly spent higher time in the target quadrant in comparison with the STZ group (p < 0.05) (Fig.4C). In this trial, platform zone crossing in the STZ group significantly decreased compared with the control and sham groups (p < 0.0001). Moreover, the STZ+Donp group had a significant difference from the STZ group (p < 0.01) and the control group (p < 0.05) (Fig.4D). Escape latency in visible platform trials was not significantly different between the groups (Fig.4E).

**Figure 4.**
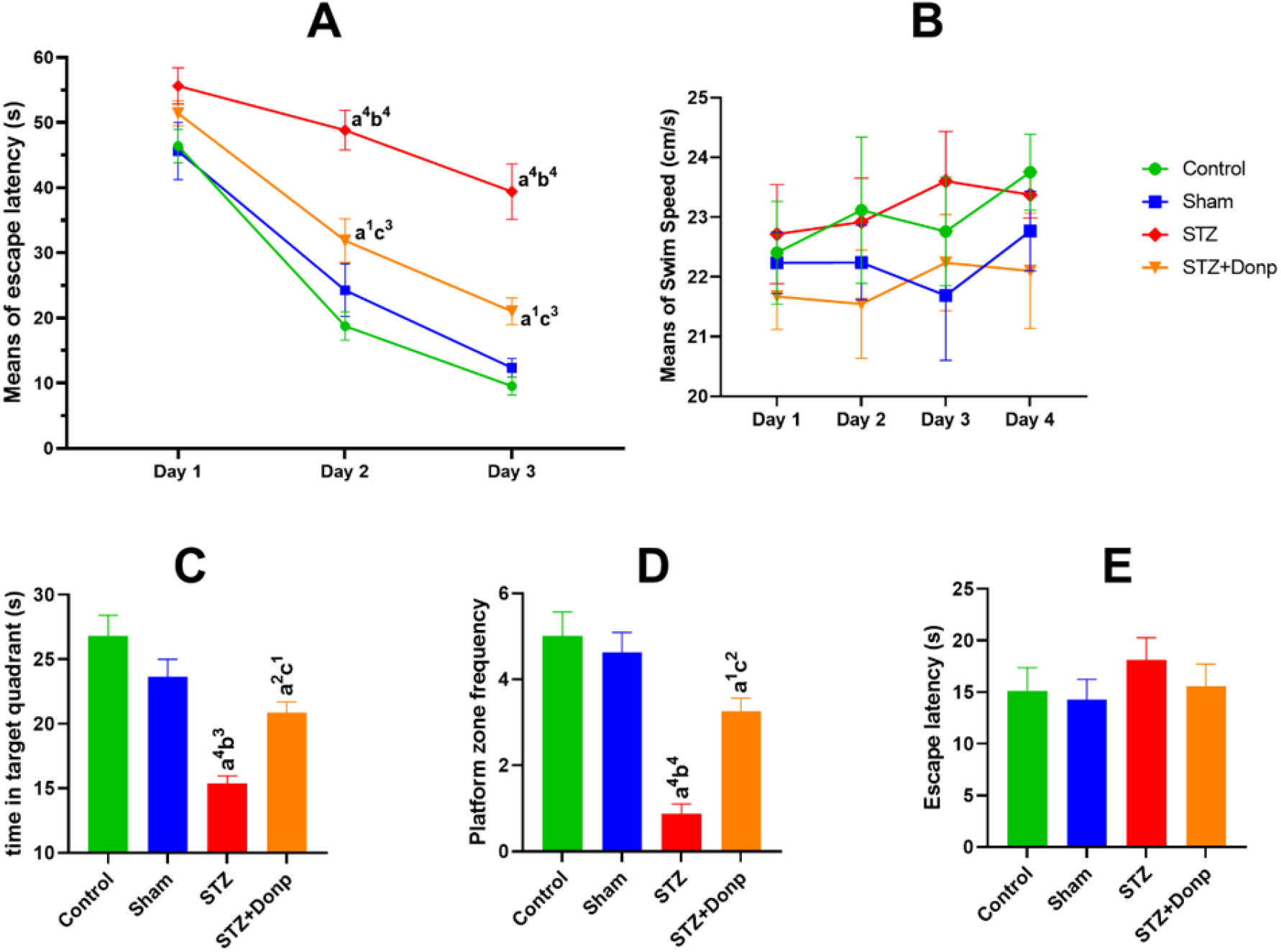
Morris Water Maze test. Data were expressed as mean ± SEM (n = 8). (A), Line chart of block mean of escape latency in trial days; (B), Line chart of velocity in trial day; (C), The target quadrant time spent in the probe trial; (D), The platform zone crossing frequency in the probe trial; (E), Escape latency in visible platform trial. Data were analyzed by two-way repeated-measures ANOVA for acquisition trial and velocity, and one-way ANOVA for probe and visible trial followed by Tukey’s multiple comparison test. (a), Compared to the control group; (b), Compared to the sham group; and (c), Compared to the STZ group; 1p < 0.05, 2p < 0.01, 3p < 0.001, 4p < 0.0001. STZ: Streptozotocin, Donp: Donepezil

### 3.2 Novel object recognition test

No differences in the spent time exploring the two identical objects during the familiarization phase were found among animals (Fig.5A). ICV-STZ injection impaired responding in the NOR test, as was indicated by failure to discriminate between familiar and novel objects during the testing phase 1h after the familiarization phase. In control and sham groups, a significant increase (p < 0.001) in exploration time towards a novel object was observed in comparison with a familiar object. Administration of donepezil improved memory in STZ -injected rats, as was shown by a significant increase (p < 0.01) in the the exmeansation time of the novel object compared with the familiar object (Fig.5B). When results were expressed as discrimination index, the STZ group had a significantly lower discrimination index compared with control and sham groups (p < 0.01, and p<0.001 respectivly), two-way ANOVA analyzed data significantly higher discrimination index in comparison with the STZ group (p < 0.05) (Fig.5C).

**Figure 5.**
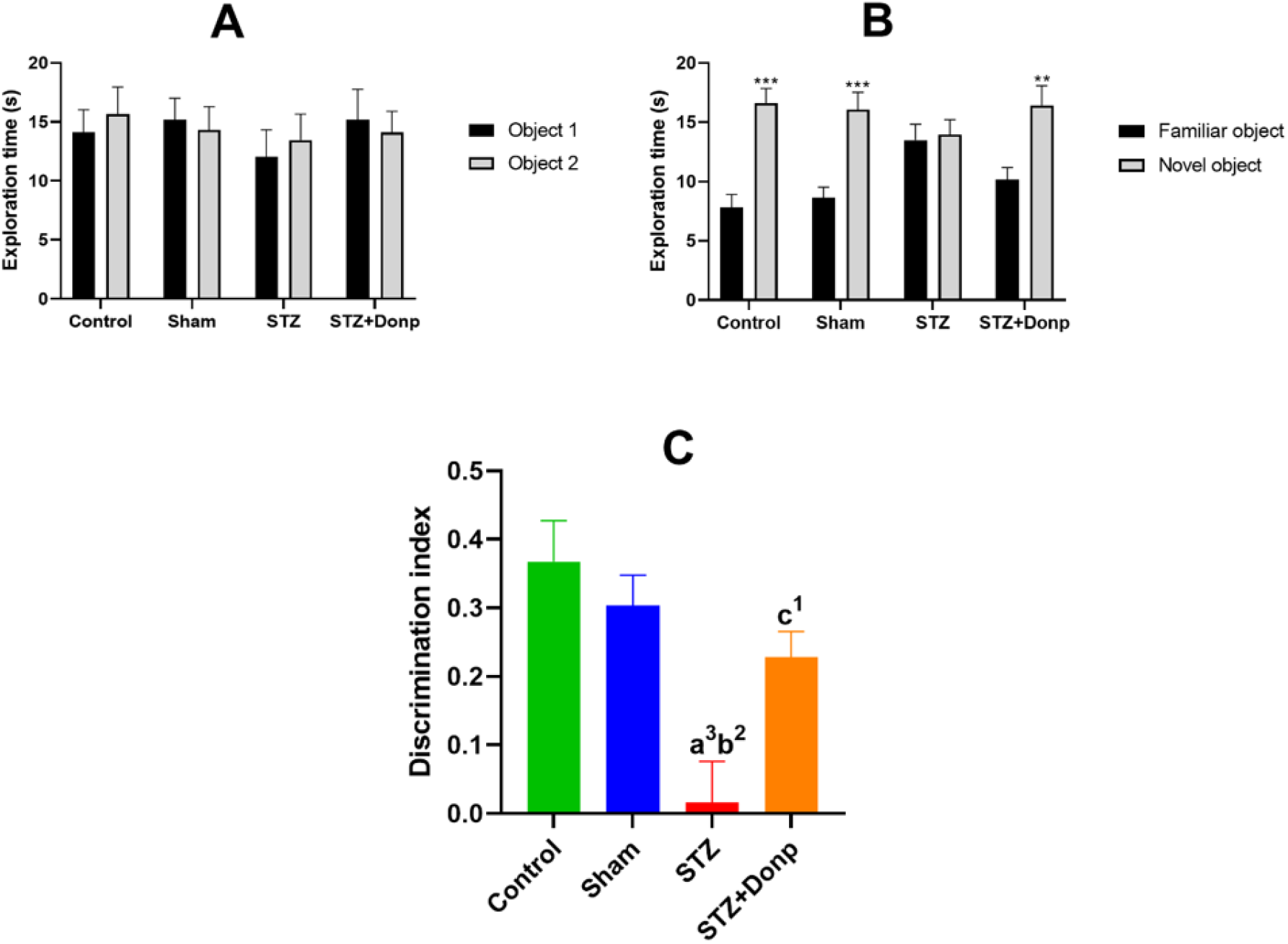
Novel object recognition test. Results were expressed as mean ± SEM (n = 8). (A), Exploration time during the familiarization phase; (B), Exploration time during the testing phase; (C), Discrimination index. Data were analyzed by paired t-test. 0.05, **p < 0.01, ***p < 0.001 compared to the exploration time of familiar objects during the testing phase of respective groups. Data were analyzed by one-way ANOVA followed by Tukey’s multiple comparison tests. (a), Compared to the control group; (b), Compared to the sham group; (c), Compared to the STZ group; ^1^p < 0.05, ^2^p < 0.01, ^3^p < 0.001. STZ: Streptozotocin, Donp: Donepezil

## 4 PHOTON EMITTING EVALUATION OF THE HIPPOCAMPUS

The mean of the total photon emission of the hippocampus during the 300 seconds was shown in Fig.6. ICV-STZ injection significantly increased hippocampus photon emission of STZ-injected rats compared with control and sham groups (p < 0.0001). The STZ+Donp rats had significantly lower hippocampus photon emission in comparison with the STZ group rats (p < 0.01).

**Figure 6.**
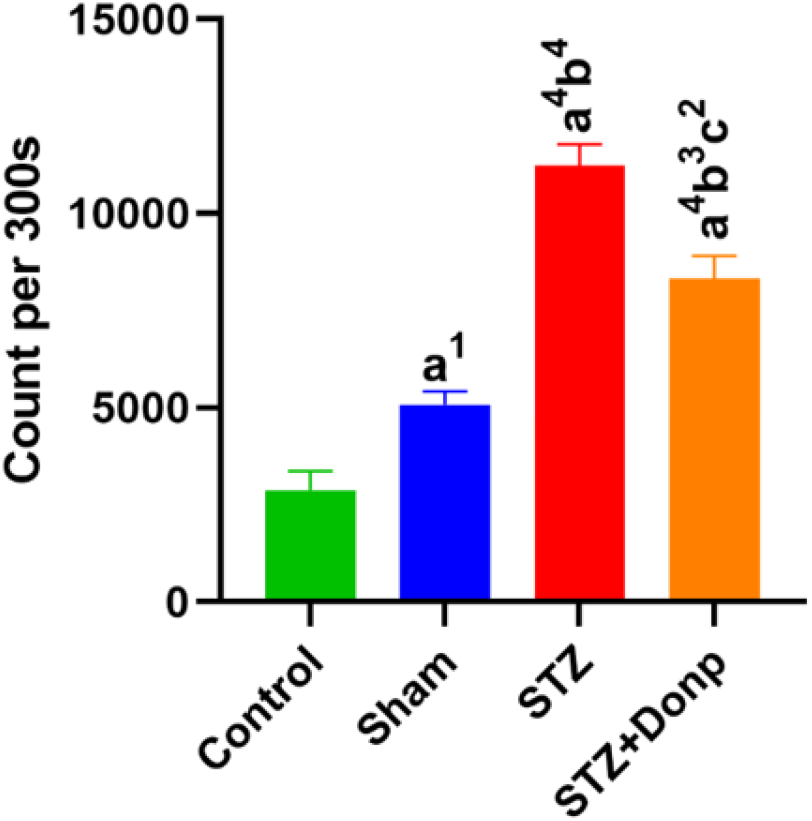
Representation of hippocampus total photon emission during the 300 seconds. The result was expressed as mean ± SEM (n = 8). Data were analyzed by one-way ANOVA followed by Tukey’s multiple comparison test. (a), Compared to the control group; (b), Compared to the sham group; (c), Compared to the STZ group; ^1^p < 0.05, ^2^p < 0.01, ^4^p < 0.0001. STZ: Streptozotocin, Donp: Donepezil

### 4.1 Hippocampus MDA concentration

MDA concentration (index of lipid peroxidation) in the STZ-injected groups was significantly higher than in control and sham groups (p < 0. 0001). Also, lipid peroxidation decreased in the STZ+Donp group (p < 0.001) compared with the STZ group but was still significant with control and sham groups (p<0.001, p<0.05, respectively) (Fig.7).

**Figure 7.**
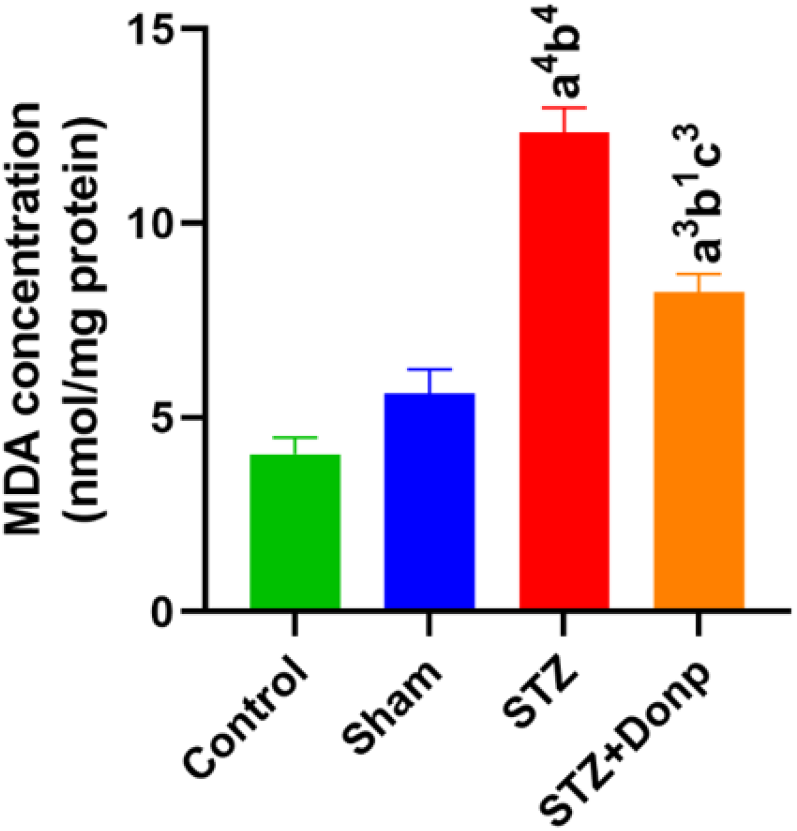
Representation of MDA concentration. Data were expressed as mean ± SEM (n = 8) and analyzed by one-way ANOVA followed by Tukey’s multiple comparison test. (a), Compared to the control group; (b), Compared to the sham group; (c), Compared to the STZ group; ^1^p < 0.05, ^3^p < 0.001, ^4^p < 0.0001. STZ, Streptozotocin; Donp, Donepezil

### 4.2 AChE activity of the hippocampus

The hippocampal AChE activity significantly (p < 0.0001) increased in the STZ-group compared with control and sham groups. In the donepezil-treated group, AChE activity of the hippocampus significantly decreased in comparison with the STZ group (p < 0.01) but had significantly higher AChE activity than in the control group (p < 0.01) (Fig.8).

**Figure 8.**
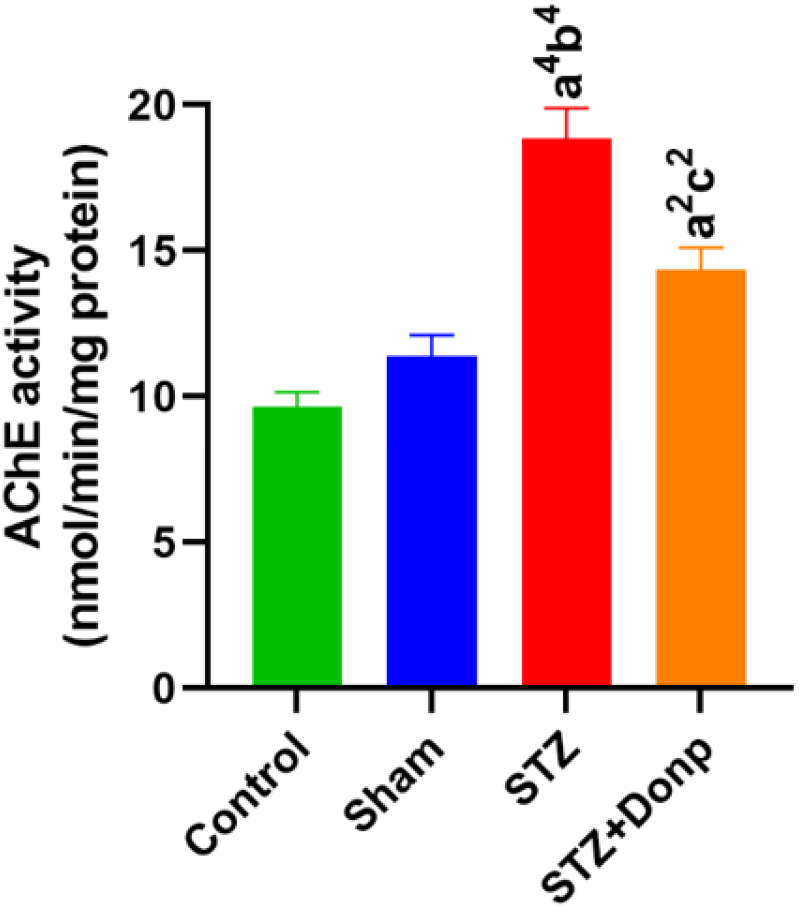
Representation of the AChE activity of the hippocampus. Data were expressed as mean ± SEM (n = 8) and analyzed by one-way ANOVA followed by Tukey’s multiple comparison test. (a), Compared to the control group; (b), Compared to the sham group; (c), Compared to the STZ group; ^1^p < 0.05, ^2^p < 0.01, ^3^p < 0.001, ^4^p < 0.0001. STZ, Streptozotocin; Donp, Donepezil.

### 4.3 Correlation matrix of UPE with MDA & AChE activity of the hippocampus

UPE had more correlation strange with MDA as was proven by Pearson correlation analysis, UPE vs. MDA (r = 0.8552); UPE vs. AChE activity (r = 0.7793). The Coefficient of determination (r2) in this analysis showed that 73% of the UPE variance could be explained by MDA concentration, while 60% of UPE variance could be explained by AChE activity (Fig.9).

**Figure 9.**
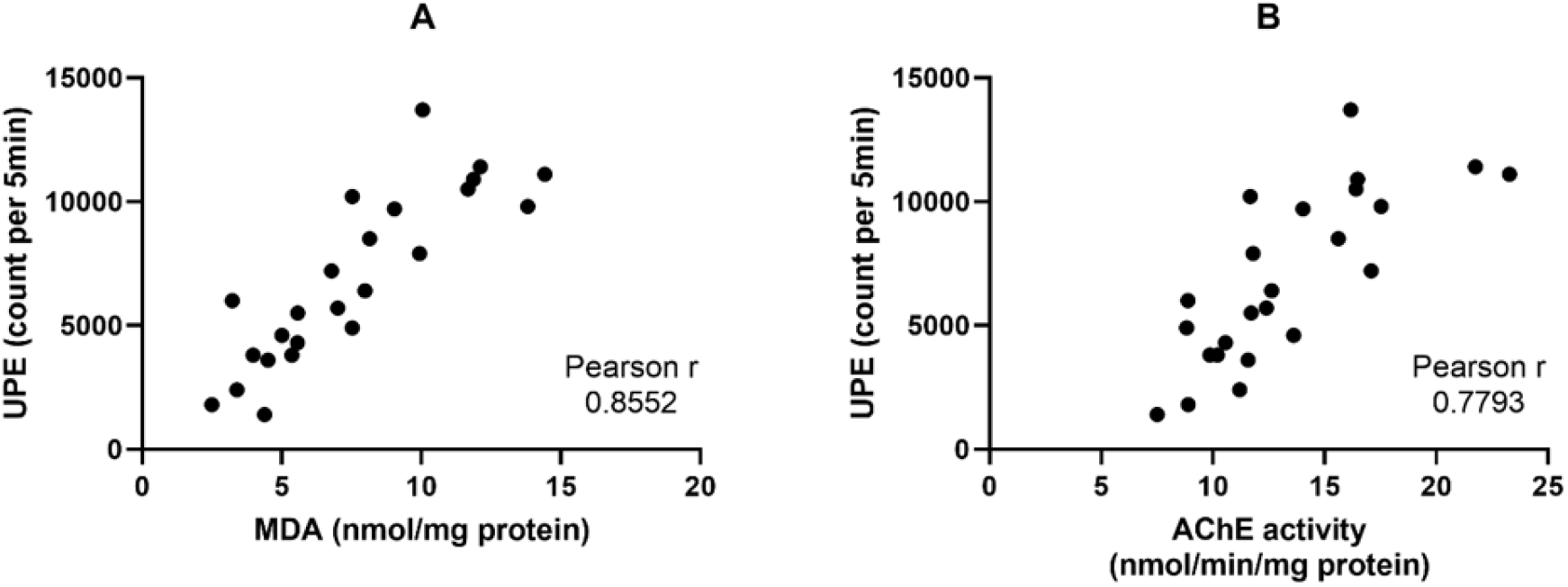
Correlation matrix of right hippocampus UPE vs. left hippocampus MDA concentration & Ache activity. (A), UPE vs. MDA; (B), UPE vs. AChE activity.

## 5 DISCUSSION

The present study investigated the effect of the most potent treatment of AD (donepezil) on the associations between the memory, intensity of UPE, oxidative stress, and AChE activity of the hippocampus for diagnosis and treatment success in ICV-STZ-induced sporadic AD models. Following ICV administration of STZ, rats exhibited memory impairment while testing for their behavioural paradigms by MWM and novel object recognition NOR tests. Previous studies regarding ICV-STZ-treated rats showed impairment in memory without any significant changes in escape latency time in MWM (40; 41) and NOR tests. Also, ICV-STZ rats had poor discrimination index and reflected no reaction to the novel object (34; 42). An anticholinesterase inhibitor (donepezil) was used as a standard treatment, and it was found that donepezil prevented ICV-STZ-induced memory impairment in all behavioral paradigms. In agreement with these results, prior studies indicated that donepezil inhibits memory deficits induced by ICV-STZ in rats (43; 44). Differences between groups in behavioural paradigms were not associated with any changes in vision and locomotor activity, as was demonstrated by no sign between groups in swim velocity and visible platform trials. The memory performance deficits induced by ICV-STZ injection may be related to various mechanisms which play an essential role in cognitive function, like disrupting the mitochondrial membrane potential and decreasing the generation of ATP (45; 46). In response to this effect, STZ induces a rise in the production of ROS and disrupts mitochondrial function to generate H2O2. When ROS are spontaneously produced during the metabolic processes, the ultra-weak Photons are emitted spontaneously from electron energy level changes in chemical reaction of electron excited species to electron ground metabolites (18). In agreement with this evidence, we detected a significant elevation in photons emitting from the hippocampus of ICV-STZ-injected rats. Kobayashi et al. found that when whole brain slices are examined under an inhibitor of the mitochondrial electron transport chain, photon emission intensity increased, indicating electron leakage from the respiratory chain (47). Biophoton emission implicates a pathophysiological state driven by the excessive production of ROS and oxidative stress. A study showed that increased UPE in rats’ brains is related to oxidative stress (48). Some reports addressed that ICV-STZ injection can lead to oxidative stress, neuroinflammation, and ROS production in rats’ hippocampus (49; 50). Consistent with these reports, we found a significant increase in MDA concentration in the hippocampus of ICV-STZ-injected rats. MDA is the index of lipid peroxidation and is known as an oxidative stress marker. It is supposed that excited species for photon emission are formed through a radical reaction with intracellular substances. Particularly regarding unsaturated fatty acids, excited species are generated as the result of a lipid peroxidation process (51). From obtained results and previous studies could be concluded that elevated biophoton emitted by the hippocampus in ICV-STZ-injected rats is linked with ROS production and MDA concentration, as was evidenced in this study by Pearson analysis of UPE vs. MDA results. A study in a mouse model of rheumatoid arthritis stated that increased UPE intensity is strongly related to metabolic processes, which may be associated with lipid oxidation and inflammatory and ROS-mediated processes. Thus, UPE may serve as a valuable tool for diagnosing chronic disease (28). Investigations in the biophoton field have considered UPE as a new potential tool for monitoring biological processes related to ROS changes, such as ROS-related diseases (52; 53) and processes pertaining to oxidative stress metabolism. UPE detection advantage is providing cost-effective spatiotemporal information without a need for invasive methods (54). The use of UPE as a non-invasive tool for different illnesses diagnosis has been proposed and discussed in different kinds of literature (55; 56). One study showed a slight increase in the UPE intensity in mice with transplanted bladder cancer, and untreated cancerous regions provided higher UPE intensity than normal regions (57). Takeda et al. evidenced changes in the UPE during the cell proliferation of human esophageal carcinoma cells (58) and tumor progression in transplanted mice (59). Boveris et al. characterized photon emission from mammalian organs (60). Evidence suggested noninvasive monitoring of oxidative metabolism and oxidative damage of living tissue using UPE (51). A document also showed that hypermetabolism induction in a rat’s brain leads to increased ultraweak photon emission as a reflection of oxidative stress (48). ROS production has been analyzed by photometry, luminometry, flow cytometry, and precipitation reaction techniques (61). All of these techniques measure at only a single time point or require chemical labels; also, these techniques are biopsy-dependent and not necessarily feasible for diagnostic purposes. While UPE is a new promising tool used to monitor real-time oxidative processes without these requirements. The utility of UPE as a tool for monitoring health and disease has been indicated in several studies (62; 63; 64). It is known that AD has a long latent period before diagnosing disease symptoms. Recent studies demonstrated that mild cognitive impairment (MCI) is common along with AD, MCI subjects exhibited significant oxidative imbalance compared with age-matched controls, and some studies have revealed that oxidative stress occurs before forming plaques and NFTs (65; 66; 67). According to obtained results and previous studies, it seems that elevated UPE of the hippocampus in ICV-STZ-injected rats is linked with ROS production and MDA concentration. It may be possible to use UPE for early detection of AD. Treatment with donepezil significantly reduced biophoton emitting in the hippocampus of ICV-STZ-injected rats. This effect seems to be caused by decreasing ROS production and oxidative stress, as was proven by the MDA concentration decrease in donepezil-treated rats compared with the STZ group. In agreement with our results, previous studies showed that treatment with donepezil had an antioxidative effect in the ICV-STZ rat model, and the treatment decreased MDA and increased GSH levels, showing the reduction of oxidative stress in the brain of ICV-STZ rats (35; 68). ICV-STZ injection in rats causes reduced energy metabolism and synthesis of acetyl CoA, ultimately resulting in cholinergic deficiency; and, thereby, memory deficit. This process is supported by reducing choline acetyltransferase (ChAT) activity (69) and increasing acetylcholinesterase (AChE) function (68) in the hippocampus of ICV-STZ-injected rats. In this line, our results manifested that AChE activity increased in ICV-STZ-injected rats. Thus, UPE may also be correlated with metabolic systems involved in neurotransmission, as evidenced by the 61% coefficient of determination (r2) between UPE and AChE activity. A study identified associations between UPE intensity and neurotransmitter metabolites (28). Donepezil, a second-generation cholinesterase inhibitor, is used therapeutically for mild to moderate dementia of AD. It inhibits AChE reversibly and non-competitively (70). Thus, donepezil treatment can decrease the hippocampus’s AChE activity in the STZ-injected rats, and this AChE metabolism reduction may be a reason for significant UPE reduction in donepezil-treated rats. But activity was not reached in the sham or control group because AChE activity was evaluated six days after the last dose.

### 5.1 Discussion of envisaged BCIPC for AD

Our compelling results calls for the development of an advanced BCI photonic chip (BCIPC) for detecting, diagnosing and classifying neurodegenerative diseases (and associated sydromes), such as in AD. The envisaged BCIPC would be minimally invasive, cheap, high-speed, scalable providing high spatiotemporal resolution of brain’s activity when compared to electrical chips, which have stronger limitations on the number of electrodes. The BCIPC would longitudinally capture brain’s UPE and appropriate data processing would complement existing neuropathological parameters to provide an enhanced clinical assessment of AD. The envisaged BCIPC is rendered in Fig. 10, and shows the possibility of connecting the BCIPC to a smart wristwatch or smartphone via its wireless connection. This technological proposal is timely, since photonic technologies are rapidly advancing. They are poised to overtake many electrical technologies due to their unique advantages, such as miniaturization, high speed, low thermal effects, and large integration capacity that allow for high yield, volume manufacturing, and lower cost. Here we provide some predictions and discuss the feasibility of the technology and its limitations for AD and related dementias. To implant a photonic chip for monitoring photon emissions from the hippocampus, a minimally invasive approach is preferred. This can be achieved through low-invasive chip transplantation surgery that involves placing the chip on the surface of the temporal skull. The temporal lobe, particularly the medial region, is closely related to memory and time episode formation, and is often affected earlier in dementia patients compared to other brain regions. Therefore, detecting UPE (ultraweak photon emissions) from the temporal lobe using a minimally invasive surface skull transplantation of a photonic chip is likely to be more feasible than monitoring hippocampal formation, which requires more invasive procedures and is more challenging to transplant. Using a photonic interferometer to distinguish wavelengths of UPEs, the efficiency decreases significantly. This low number of photons is still interpretable since the chip will be fitted with single-photon detectors that receive a relatively high number of photons compared to the quantum limit (i.e., 10-10^3^ photons/sec). Moreover, the signal will be further discriminated from noise (e.g., dark noise and shot noise) via appropriate machine-learning training data (for a detailed discussion we refer the reader to our previously published work (3)). The integration time of the detector is estimated to be on the order of nanoseconds. Generally, biological systems exhibit UPE spectra from the near-ultraviolet and extend to the range 700-1000 nm, an optimal range for our photonic detector. Similar results may be achieved in the ultraviolet and visible spectra. These combined advantages enable for the future development of a ‘‘brain photonics” detection device to passively image the brain’s spontaneous (Fig.10).

**Figure 10.**
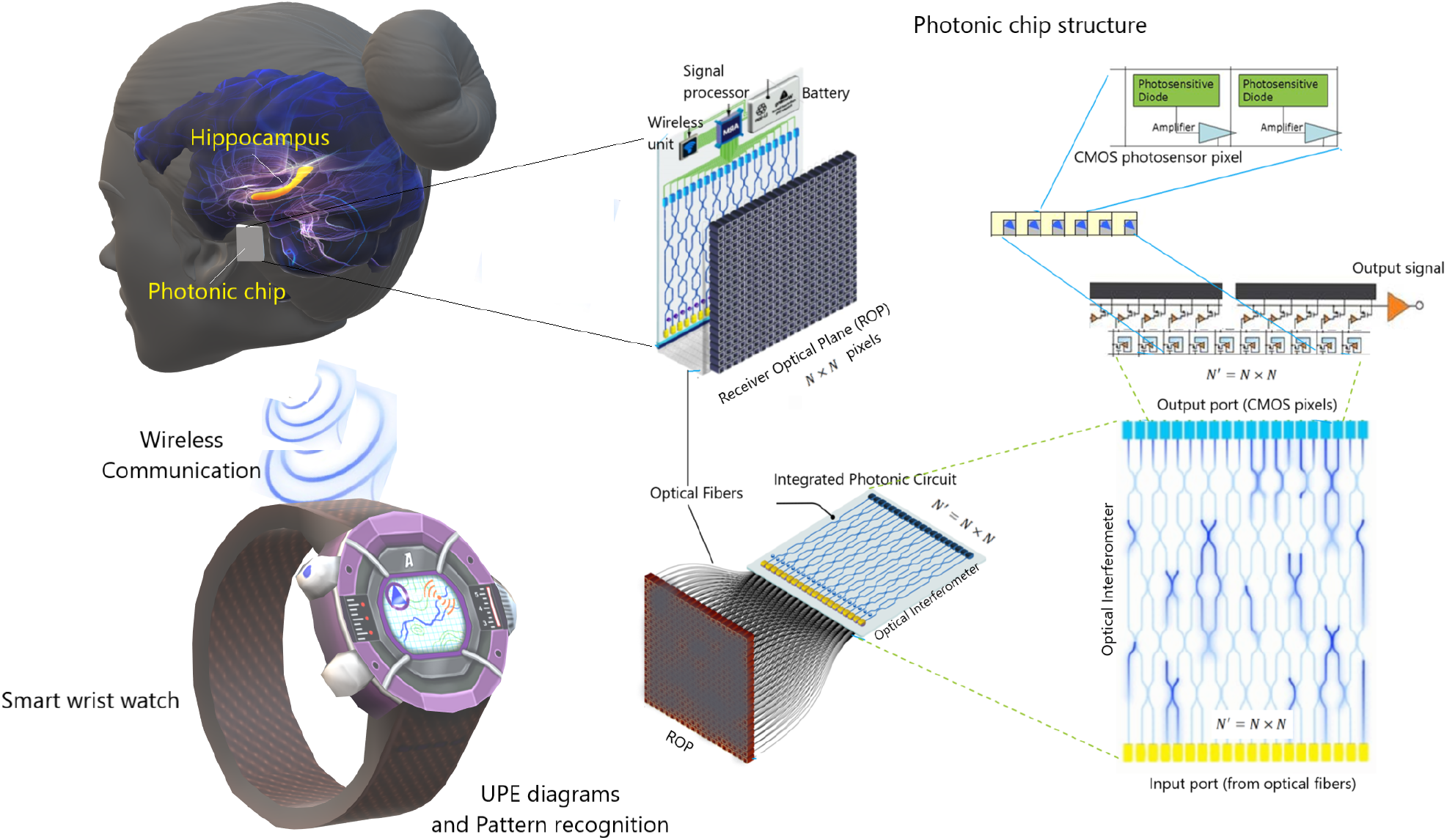
A schematic representation for futuristic monitoring hippocampus UPE variations and pattern recognition that may help better diagnosis of Alzheimer’s disease in the short- and long-term. The photonic chip can be connected to a smart wristwatch and monitor the brain state continuously. The figure is an update from the previously published work(3). For more details about the chip structure, see Ref. (3).

## 6 CONCLUSION

Treatment with donepezil improved spatial and recognition memory deficits via oxidative stress regulation and AChE inhibition in ICV-STZ-injected rats. Correlations with UPE led us to the conclusion that hippocampus UPE is associated with the redox state of the tissue. Since oxidative stress is one of the primary patho-neurophysiological signatures in the progression of AD, then it stands to reason that UPE detection of the brain will provide complementary clinical parameters for AD screening and diagnosis. Moreover, UPE could be used to monitor recovery from neurodegenerative diseases upon suitable future therapeutic treatments, as suggested by our experiment involving donepezil which decreases the patho-neurophysiological signatures and in a correlated way with UPE. In the cytosol of cholinergic presynaptic neurons, Ach neurotransmitter is produced from choline and acetyl-coenzyme A (acetyl-CoA) by means of choline acetyltransferase (ChAT). Acetyl-CoA is a key energy precursor intermediate in all cells of our body. Acetyl-CoA is almost entirely synthesized in the brain by the pyruvate dehydrogenase multi-enzyme complex (PDHC), which supplies 97 percent of the energy (71). Current AD therapy is mainly based on inhibitors of AChE by AChE inhibitors such as donepezil, rivastigmine, memantine, and galantamine, which enhance cholinergic transmission (72). The use of donepezil is particularly promising, as it also has potent anti-inflammatory effects, inhibits neuronal death and cognitive decline, and reduces pro-inflammatory gene expression (72). In the AD model, ICV-STZ injection induces AD disease-like symptoms that look like biomolecular, pathological, and behavioural features of AD (73; 74). ICV-STZ produces mitochondrial dysfunctions such as anomalous morphology, decreased ATP synthesis, and increased ROS generation (74; 75). This may explain why ICV-STZ injection significantly increased UPE and MDA concentrations in the hippocampus compared to the sham and control groups. Furthermore, donepezil inhibits AChE, which catalyzes the breakdown of acetylcholine (and some other choline esters that function as neurotransmitters).

This may reduce mitochondrial acetyl-CoA production, i.e., mitochondria produce less acetyl-CoA, which changes the mitochondrial redox state and various mitochondrial mechanisms. As a result, these processes reduce UPE. However, the mitochondrion is the major redox, cellular signalling, and energetic hub of cells and neurons (76). In our experiments, treatment with donepezil improved spatial and recognition memory deficits via oxidative stress regulation and AChE inhibition in ICV-STZ-injected rats. Therefore, based on the obtained results, it could be concluded that hippocampus UPE is associated with the redox state of the tissue. Pearson correlation analysis revealed that 73% of the UPE variance could be explained by MDA concentration, while 60% of the UPE variance could be explained by AChE activity of the hippocampus. In addition, studies have suggested that mitochondria can have key roles in neurogenesis (77; 78; 79). Increasing evidence suggests that the dysfunction of cellular organelles, particularly mitochondria, has important roles in neurodegenerative disorders (79). These above-mentioned results and facts support the hypothesis that perturbed redox (oxidative stress) and mitochondrial mechanisms may play key roles in the development of AD. Thus, the UPE detection of the brain may be useful for a better understanding of the development of AD, its diagnosis, and the development of possible drugs. Therefore, it is possible that a photonic chip that can efficiently detect biophotons from the hippocampus could be a tool for the diagnosis or monitoring of AD. Biophoton emissions from the brain have been suggested as a potential diagnostic marker for various neurological disorders, and here we have shown that it can include AD, in which detecting changes in biophoton emissions from the hippocampus could help monitor the progression of AD. Since the results for UPE measurement are obtained with PMT in our experiments, we suggest that the photonic chip with a similar quantum efficiency as PMT can be a helpful tool for the diagnosis of AD. The CMOS quantum efficiency is about 75%, which is about three times higher than PMTs with a quantum efficiency of about 20-25%. According to the estimations in Ref. (3), the amount of total photon loss from the receiver optical plane (ROP) to the output of the optical interferometer (OI) is about 50%. The QE of CMOS at the output of the OI is estimated to be 25% in body temperature under the implant conditions to have a final SNR of about 2. However, still, further research is needed to validate the use of biophoton emissions as a diagnostic tool and determine the specific changes in biophoton emissions associated with AD, while we have now a piece of evidence to be hopeful for designing new and cheaper methods for AD diagnosis.

## AUTHOR CONTRIBUTIONS

Conceptualization: TE, AZ, FD, VS; Data curation: NS, TE, MKG; Formal analysis: TE, VS, MKG; Funding acquisition: TE, VS; Methodology: TE, NS, AZ, FD, MKG; Project administration: TE, VS; Resources: TE, AZ; Supervision: TE, VS, DO; Discussion: NS, TE, VS, MKG; Hypothesis: NS, TE, VS; Writing an original draft: NS, TE, VS, MKG; Writing a review and editing: TE, VS, NC, IB, SR, and DO.

## ACKNOWLEDGMENTS

VS and DO are grateful for fruitful discussions with Christoph Simon and Hadi Zadeh-Haghighi. SR acknowledges support from Ikerbasque (The Basque Foundation for Science), the Basque Government through the BERC 2022-2025 program and by the Ministry of Science and Innovation: BCAM Severo Ochoa accreditation CEX2021-001142-S / MICIN / AEI / 10.13039/501100011033 and through project RTI2018-093860-B-C21 funded by (AEI/FEDER, UE) and acronym “MathNEURO”.

## DATA AVAILABILITY STATEMENT

The datasets [GENERATED/ANALYZED] for this study can be found in the [NAME OF REPOSITORY] [LINK].

